# Recapitulation of human germline coding variation in an ultra-mutated infant leukemia

**DOI:** 10.1101/248690

**Authors:** Alexander M Gout, Rishi S Kotecha, Parwinder Kaur, Ana Abad, Bree Foley, Kim W Carter, Catherine H Cole, Charles S Bond, Ursula R Kees, Jason Waithman, Mark N Cruickshank

## Abstract

*Mixed lineage leukemia/Histone-lysine N-methyltransferase 2A* gene rearrangements occur in 80% of infant acute lymphoblastic leukemia, but the role of cooperating events is unknown. While infant leukemias typically carry few somatic lesions, we identified a case with over 100 somatic point mutations per megabase and here report unique molecular features of this peculiar case. The patient presented at 82 days of age, one of the earliest manifestations of cancer hypermutation recorded. The transcriptional profile showed global similarities to canonical cases. Coding lesions were predominantly clonal and almost entirely targeting alleles reported in human genetic variation databases with a notable exception in the mismatch repair gene, *MSH2*. There were no rare germline alleles or somatic mutations affecting proof-reading polymerase genes *POLE* or *POLD1*, however there was a predicted damaging mutation in the error prone replicative polymerase, *POLK*. The patient’s diagnostic leukemia transcriptome was depleted of rare and low-frequency germline alleles due to loss-of-heterozygosity, while somatic point mutations targeted low-frequency and common human alleles in proportions that offset this discrepancy. Somatic signatures of ultra-mutations were highly correlated with germline single nucleotide polymorphic sites indicating a common role for 5-methylcytosine deamination, DNA mismatch repair and DNA adducts. Thus, we document a novel somatic ultra-mutational signature that recapitulates population-scale human genome variation with unprecedented precision.

## Author Summary

### Retraction

This article should not be cited. We are currently investigating alternative constitutional patient DNA from this patient. She was treated upfront by stem cell transplantation. The timing of remission sample collection with respect to the transplantation requires confirmation. It is plausible the remission sample sequenced includes transplanted cells. We apologise for the inconvenience.

The mutational burden of human cancer varies widely. Infant cancers typically have few somatic alterations. In contrast, some tumours including 16% of adult cases and 6% of childhood cases, display an elevated mutation rate due to either environmental exposures or genetics. The most highly mutated human cancers (“ultra-mutated”) typically involve mutations and/or inherited defects in a DNA mismatch repair gene (commonly *MSH2, MSH6, MLH1* or *PMS1)* and a proof-reading DNA polymerase gene (either *POLE* or *POLD1)*. Infant leukemias typically carry few somatic lesions, and as a step to characterise the combined germline and somatic variations we analysed the coding sequences and transcriptomes of six patients’ remission-diagnosis samples. Most infant samples showed a “silent” genome with few somatic lesions. However, we observed a case diagnosed at 82 days of age with an ultra-mutated phenotype - the earliest manifestation on record. The sample had a mutation in the DNA-mismatch repair gene, *MSH2* but there were no co-mutations affecting proof-reading polymerase genes *POLE* or *POLD1*. An alternative secondary hit, was observed in the replicative error-prone DNA polymerase *POLK*. Remarkably, the somatic ultra-mutations in this patient precisely targeted rare and common human germline alleles effectively causing a widespread “shuffling” of germline alleles without increasing the rare allele burden. Thus, the ultra-mutations in this sample mirror important aspects of human genome evolution and represent a novel somatic mutational mechanism.

## Introduction

*Histone-lysine N-methyltransferase 2A (KMT2A)* (also known as *Mixed lineage leukemia)* gene rearrangements *(KMT2A-R)* at 11q23 occur in infant, pediatric, adult and therapy-induced acute leukemias and are associated with a poor prognosis. *KMT2A-R* occurs in around 80% of infant acute lymphoblastic leukemia (ALL) and 35–50% of infant acute myeloid leukemia (AML) involving more than 60 distinct partner genes ^41^. *KMT2A-R* infant ALL (iALL) genomes have an exceedingly low somatic mutation burden, with recurring mutations in *RAS-PI3K* complex genes. Compared to wild-type KMT2A, the fusion oncoprotein has altered histone-methyltransferase activity, causing epigenetic and transcriptional deregulation. While KMT2A-oncoproteins are strong drivers of leukemogenesis, several lines of evidence suggest that *KMT2A-R* alone is insufficient for tumor initiation. For example, there is significant variability in latency of disease onset in experimental models of KMT2A-fusions ^6^, ^14^ and in patients with *KMT2A-R* detected in neonatal blood spots ^24^. Furthermore, *KMT2A-R* acute leukemias ^5^ in children are enriched for mutations affecting epigenetic regulatory genes and on average harbor more somatic lesions than infants. These observations have prompted speculation that iALL is a distinct developmental and genetic entity and pre-leukemic transformation occurs *in utero*, targeting a cell of origin with a genetic/epigenetic permissive predisposition ^48^.

As a step towards defining cooperating events we performed next-generation sequencing on a *KMT2A-R* iALL patient cohort and integrated the data with a larger set of acute leukemias. We investigated the combined germline and somatic mutation profiles, identifying a highly mutated specimen from a patient diagnosed at 82 days of age. Here we report sequence analysis of this unique leukemia in relation to its canonical *KMT2A-R* iALL counterparts, revealing common mechanisms driving leukemia ultra-mutation and population-scale human genome variation.

## Results

### Characteristics of KMT2A-R iALL cohort and RNA-seq variant detection pipeline

We reported previously *KMT2A-R* fusion transcripts for 5/10 cases ^18^, and of the remaining five *KMT2A-R* iALL we detected *KMT2A-MLLT3* (P401, P438), *KMT2A-AFF1* (P848), *KMT2A-MLLT1* (P706) and *KMT2A-EPS15* (P809). KMT2A-fusion transcripts were therefore detected in all samples and reciprocal fusion sequences detected in 5/10 samples (Table 1). To determine the validity of our RNA-seq based pipeline for detecting rare and common alleles, we examined the concordance of single nucleotide variants (SNVs) called from RNA-seq and whole exome sequencing (WES) data. Across seven samples with matched WES and RNA-seq data, we identified a total of 54,792 point mutations from RNA-seq reads of which 166 were absent from matched WES (coverage>20 reads), representing a discordance rate of 0.3% (Figure 1A; Supplementary Table S2). In one sample, P706 there were two events called with RNA-seq data that were not identified in matched WES but with evidence extracted using the Integrated Genome Browser at previously reported somatic mutation sites in *FLT3* (encoding FLT3 p.V491L ^25^; 200/1,719 alternate RNA-seq reads; 15/214 alternate WES reads) and *KRAS* (encoding KRAS p.A146T ^11^; 24/159 alternate RNA-seq reads; 6/51 alternate WES reads). Furthermore, analysis of 117 discordant events with coverage > 50 reads, revealed 76 recurrences in two or more samples (64%) at 26 unique sites, however none of these sites have been validated as somatic mutations in cancer to date in the Catalog of Somatic Mutations in Cancer (COSMIC ^22^) or The Pediatric Cancer Genome Project (PCGP ^20^) databases. These results indicate a high rate of reproducibility of the RNA-seq variant detection pipeline and suggest that RNA-DNA differences mostly comprise artifacts and/or novel RNA-editing events ^43^ and a minority of putative somatic alterations exclusively detected by RNA-seq.

**Figure 1:** Characterization of germline and somatic point mutations identifies an ultra-mutated infant leukemia case. (A) Overlap of variants identified from diagnostic leukemia specimen by RNA-seq and whole exome sequencing (WES). (B) Overlap of variants called in diagnostic RNA-seq and remission bone marrow WES. (C) Overlap of variants called in diagnostic and remission WES. Barplots colored showing overlap binned by derived allele frequency as follows: present with AF<0.5% in ExAC (green), present with AF 0.5–1% in ExAC (yellow), present with AF>1% in ExAC (purple), absent from the ExAC (orange), non-overlapping variants are colored to show sites with coverage <20 reads (grey) or coverage >20 reads (black) in the comparator. (D) Scatterplots of diagnostic leukemia (x-axis) versus remission (y-axis) variant allele fraction (VAF) from WES for coding variants detected in diagnostic leukemia RNA-seq. (E) Dendrogram showing sample clustering computed with WES variants from primary diagnosis/remission samples and cell lines (PER-784A, PER-826A).

### Identification of a highly mutated infant leukemia

Expressed somatic mutations were defined as variants detected in both diagnostic leukemia RNA-seq and WES, but absent in remission WES at sites with >20 reads (Figure 1). Most patients (5/6) showed the expected “silent genomic landscape” (Supplementary Table S3) except for patient P337 (Figure 1A-D; Supplementary Table S4). Three patients expressed RAS-family hot-spot somatic mutations (P848 and P706: KRAS p.A146T and P438: NRAS.p.G12S). We detected no expressed somatic missense mutations in patients P399 and P401. The pair of monozygotic twins (P809 and P810) lacked private expressed mutations but we detected two shared expressed missense mutations clustered in the *PER3* gene (encoding PER3.p.M1006R and PER3.p.K1007E). These results are consistent with recurring *RAS-*pathway mutations and low mutation rate reported previously ^5^, ^12^. However, in patient P337 RNA-seq reads we detected 1,420 expressed somatic mutations (Supplementary Table S4). We previously reported a complex *KMT2A* translocation involving 2q37, 19p13.3 and 11q23 ^26^ in the highly mutated leukemia sample and established two cell lines from the diagnostic sample ^18^ which were also subject to WES. Hierarchical clustering of variant data, demonstrated co-segregation of the highly mutated diagnostic specimen with matched cell lines (Figure 1E). The corresponding remission sample forms a sub-cluster indicating its relatedness to these samples, however, it was more distant from its corresponding diagnostic sample compared to canonical cases. We note that samples from monozygotic twins (P809 and P810) formed a distinct group, with individual-specific diagnosis/remission paired samples forming sub-clusters. Altogether, these results verify the common origin of RNA-seq and diagnostic/remission WES data ruling out sample contamination or mislabelling.

### Characterisation and clonal analysis of somatic lesions

Somatic lesions were further evaluated in six cases by comparing diagnostic and remission WES. Two different variant allele fraction (VAF)-cut-offs, either VAF>0.1 or VAF>0.3, were chosen to enrich for sub-clonal and clonal events respectively. There were on average ~82.2 (range 69–95) sub-clonal and ~3.4 clonal (range 3–4) somatic point mutations in canonical *KMT2A-R* iALL exomes, confirming previous studies of a low somatic mutation rate in the dominant clone (Table 1). In contrast, there were 198 sub-clonal and 5,054 clonal somatic point mutations in the highly-mutated sample, equating to a rate of 139 mut/Mb, classifying this sample as ultra-mutated. There were also ~40-fold increased number of indels in the dominant clone of the ultra-mutated sample (n=260) compared to typical cases (average = 6.6; range: 3–9) including 13 predicted to cause a frame-shift protein-coding truncation (Table 1; Supplementary Table S5). There were 2,420 exonic substitutions encoding 1,109 missense alleles and 12 stop-gain alleles (Supplementary Table S5). We also noted that the ultra-mutated case carried an excess of loss-of-heterozygosity (LOH) positions, with 6,555 high confidence sites; vastly more than typical cases which had a mean of 57.2 LOH positions (range 28–100) (Figure 2A-B; Table 1).

**Figure 2:** Clonal analysis of somatic ultra-mutations. Histograms showing the VAF density distribution of somatic mutations (left, x-axis: dark shaded bars) or LOH-sites (right; x-axis: light shaded bars) in diagnostic leukemia WES from canonical cases (A) and the ultra-mutated case (B). (C) Illustration of cell lines derived from ultra-mutated specimen and histograms displaying the VAF density distribution of somatic mutations (left, x-axis: dark shaded bars) or LOH-sites (right; x-axis: light shaded bars) in cell lines (PER-784A: red; PER-826A: green). (D) Venn diagram displaying the overlap in somatic mutations (left) and LOH-sites (right) identified by *VarScan2* comparing the primary diagnostic specimen and derived cell lines (PER-784A: red; PER-826A: green) to the primary remission sample. (E) Somatic mutation and (F) LOH coding sites showing total read coverage (left y-axis) and VAF (right y-axis) from primary specimens (left panels) and cell lines (right panels) ordered by decreasing VAF in the diagnostic sample (x-axis).

WES was performed on two independent cell lines derived from the ultra-mutated diagnostic specimen, PER-784A and PER-826A, to characterize clonal somatic events. (Figure 2C,D). Most somatic and LOH-events detected in the primary ultra-mutated sample were also called in both, PER-784A and PER-826A cell lines using P337 remission WES for all comparisons (Figure 2D). These data suggest that the somatic lesions are largely clonal in agreement with the VAF-distributions (Figure 2B,D). In total, 4,754/5,054 (94%) somatic events (Supplementary Table S6) and 6,150/6,575 (93%) LOH-events (Supplementary Table S7) were called in the derived cell lines. Moreover, there were relatively few discordant sites when directly comparing cell lines with the primary diagnostic sample (PER-784A: 34 discordant sites; PER-826A: 44 discordant sites; Supplementary Table S6). We further investigated evidence of clonal somatic lesions in cell lines using an alternative approach, by directly extracting quality filtered WES reads mapping to coding somatic point mutations (n=2,411; Q>50 in diagnosis WES; Figure 2E) and LOH-sites (n=3,117; Q>50 in remission WES; Figure 2F; Supplementary Table S8). All coding clonal somatic mutations were supported by reads in both cell lines of which 121 (5%) were homozygous (VAF>0.95 in cell line and VAF>0.9 in primary sample). Most LOH-sites were evidenced by reference alleles (VAF<10%; 2,179/3,117) in cell lines, with ~30% of LOH-sites evidenced by alternate alleles (VAF> 90%; 926/3,117) and only 12 sites (0.3%) with both reference and alternate alleles (VAF range 10–90%).

We performed copy number analysis of the ultra-mutated sample to map LOH-sites to chromosomal segments with evidence of deletions. Most genomic regions were called as diploid with limited chromosomal copy loss detected (Supplementary Figure S1). There was a total of 382/6,555 (5.8%) LOH-sites mapping to homozygous deletions, suggesting only a minority of the identified LOH-sites likely arose through chromosome segment deletion. Infant *KMT2A-R* ALL typically carry a small number of copy-number alterations which may coincide with LOH-sites. We determined the distance of LOH-sites, somatic mutations and germline variants (called by *VarScan2)* to the nearest homozygous SNV in each leukemia diagnostic sample finding that in canonical cases LOH-sites tend to be located closer to homozygous sites than either germline or somatic variants (Supplementary Figure S1C; left panel) and show a unique bimodal distribution (Supplementary Figure S1C; right panel). In contrast the average distance of LOH-sites to homozygous SNVs in the ultra-mutated sample was similar to that of germline variants and somatic mutations with each displaying a similar distribution (Supplementary Figure S1C). Thus, most LOH-sites in the ultra-mutated specimen do not show evidence of localising within deleted or copy-neutral LOH-chromosomal segments.

In summary, these data demonstrate that somatic lesions in the primary sample were predominantly clonal and LOH-sites appear to comprise a large proportion of somatic substitutions rather than chromosomal alterations. We also note relatively few coding sequence changes in derived cell lines compared to the dominant diagnostic leukemic clone, both which are capable of long-term propagation. These results could reflect loss of the hypermutator phenotype during *in vitro* culture or in diagnostic leukemia cells.

### Signatures of somatic mutations and LOH-sites correlate with profiles of germline alleles

We next investigated the sequence features of somatic and LOH substitutions in the ultra-mutated sample (Figure 3). We hypothesised that LOH-sites and somatic mutations may show common feature biases depending on the context of alteration. Somatic events were evaluated separately that induce a heterozygous mutation (n=4,773) or homozygous mutation (n=281), and LOH causing reference→alternative changes (n=2,057) were evaluated separately to alternative→reference changes (n=4,453). Since we noted an overlap in the somatic mutations of the ultra-mutated sample with variants in the Exome aggregation consortium database (ExAC ^35^) (Figure 1A-C), we first examined the distribution of these sites with respect to the derived allele frequency in ExAC. We observe that each of these categories of sequence alterations in the ultra-mutated sample were distributed across the range of population allele-frequencies (Figure 3A,B). While the distributions of LOH and somatic sites were similar, somatic heterozygous changes were relatively more frequent at higher frequency alleles as were LOH reference→alternate alleles.

**Figure 3:** Germline-like somatic signatures of ultra-mutations. Histograms showing the number (A-B) or frequency (C-D) of events overlapping the spectrum of ExAC derived allele frequency bins with 5% increments. (A) Somatic mutations called by *VarScan2* grouping homozygous and heterozygous events separately. (B) Loss-of-heterozygosity sites called by *VarScan2* grouping reference→alternative and alterantive→reference sites separately. (C-D) All quality filtered coding variants from each of the canonical patients or P337 ultra-mutated specimen called by GATK grouped by single nucleotide polymorphisms (SNPs) that were homozygous and heterozygous events from remission (R) or diagnostic sample (D). (E) Normalized fraction of variants within unique trinucleotide context (ordered alphabetically on the x-axis) for *VarScan2* calls (left panel; as shown for panels a-b), or remission variants (right panel; as shown for panel C). (F) Somatic signatures identified by *deconstructSigs* for categories of mutations/SNPs (displayed in panel E) showing their relative proportions (bar plot) and pair-wise Pearson’s correlation coefficients (heatmap), ordered by unsupervised hierarchical clustering.

Somatic signatures of ultra-mutations were investigated in comparison to heterozygous and homozygous single nucleotide polymorphisms (SNPs) in patient remission samples. Heterozygous positions were defined by VAF<0.9 yielding 51,347 heterozygous and 31,237 homozygous SNPs in canonical remission samples and 10,681 heterozygous and 6 617 homozygous SNPs in the ultra-mutated remission sample. Analysis of WES remission variants from canonical patients and the ultra-mutated sample revealed similar ExAC allele frequency distributions with distinctions between heterozygous and homozygous alleles (Figure 3C,D). The overall enrichment of each category of somatic substitution and LOH across the 96 possible trinucleotide contexts resembled patient remission SNPs with elevated proportions of C>T transitions at CpG motifs (Figure 3E).

Pre-defined signatures were used to infer mutational profiles revealing Signature 1A (associated with spontaneous deamination of 5-methylcytosine), Signature 12 (associated with unknown process with strand-bias for T→C substitutions) and Signature 20 (associated with defective DNA mismatch repair, MMR) as enriched in each of the mutation/variant sets, however with distinctive proportions (Figure 3F). Heterozygous somatic mutations and LOH alternative→reference sites were additionally enriched for Signature 1B (variant of Signature 1A) and showed highly correlated profiles (Pearson’s correlation 0.919). LOH reference→alternative sites were additionally enriched for Signature 5 (associated with unknown process with strand-bias for T→C substitutions) and were highly correlated with homozygous somatic mutation profiles (0.912) and to profiles of patient heterozygous positions (P337: 0.935; canonical: 0.926). Homozygous somatic mutations also displayed similarities to heterozygous patient SNPs (P337: 0.981; canonical: 0.985). Overall the somatic mutational profiles indicate highly related patterns of both somatic ultra-mutations and LOH-sites. Notably LOH alternative→reference sites showed unique and overlapping features to somatic heterozygous mutations. We also observe underlying similarities in the profiles of somatic substitutions and germline alleles.

### Ultra-mutation distribution recapitulates common and rare germline variation

We noted a similar spectrum of rare and common alleles in the ultra-mutated leukemia transcriptome compared to typical *KMT2A-R* iALL cases; but there were comparatively fewer rare/low-frequency alleles shared with the remission sample (Figure 1A-C; ExAC<0.5%). We further explored this trend to disentangle the contribution of somatic acquired alleles and LOH-sites. Analysis of the remission and diagnostic samples from this patient in isolation, revealed a similar proportion of rare/low-frequency coding alleles detected by WES and proportionally were within the same range as canonical counterparts (Figure 4A). However, when directly comparing the remission and diagnostic WES, we noted a modest depletion of germline rare/low-frequency coding SNV alleles defined by *VarScan2* (Figure 4A; 2.9% rare alleles in P337 compared to 3.5±0.2% in canonical cases) coincident with an increased proportion of somatic acquired rare/low-frequency coding SNV alleles compared to LOH-sites. Therefore, when considering the total pool of SNVs in the leukemic coding genome, the somatic gains and losses of rare/low-frequency coding SNV alleles were in proportions that resulted in a similar distribution as canonical cases (Figure 4A). We extended this observation by investigating the rare and common allele abundance of expressed germline-encoded variants and somatic mutations in the ultra-mutated leukemic sample compared with a larger cohort of *KMT2A-R* iALL transcriptomes (Figure 4B-C). Most typical *KMT2A-R* iALL diagnostic specimens expressed similar numbers of rare and low-frequency expressed SNVs, except for 4/32 cases sequenced by St Jude investigators, which had an elevated number of expressed rare and low-frequency ExAC alleles (Figure 4B). Excluding these samples, typical *KMT2A-R* iALL patients expressed on average 42.3 ±10.4 (standard deviation) variants that were absent from ExAC and 212 ± 32.5 variants with an allele frequency of <0.5% in ExAC. The total number of expressed SNVs in the ultra-mutated sample was slightly lower than typical cases (38 absent from ExAC and 174 AF <0.5% in ExAC), comprised of approximately half germline (17/38 and 88/174 respectively) and half somatic acquired alleles. Overall, the ultra-mutated transcriptome had the lowest prevalence of germline-encoded rare and low-frequency alleles among the larger sample size (Table 2), however, somatic mutations were targeted to rare and common human alleles in proportion that off-set this difference (Figure 4B-C).

**Figure 4:** Ultra-mutations target human alleles including a rare *POLK* allele together with a novel *MSH2* allele. (A) Proportion of coding single nucleotide variants in diagnostic (D) and remission (R) whole exome sequencing (WES) data binned by derived allele frequency as follows: present with AF<0.5% in ExAC (green), present with AF 0.5–1% in ExAC (yellow), present with AF>1% in ExAC (purple), absent from the ExAC (orange). Grey shading denotes data from the ultra-mutated specimen. The distribution of variants called as germline, somatic or LOH by *VarScan2* are displayed for comparison. (B) Number of variants (y-axis) called from RNA-seq reads from St Jude and Perth cohorts (open circles) are plotted with the total variants detected in the ultra-mutated sample (shaded circles), germline-encoded variants (shaded triangle pointing up) and somatic point mutations (shaded triangle pointing down) binned according to the allele count in ExAC database (red: not in ExAC; green < 0.5%; yellow: 0.5–1% and blue > 0.5%). (C) Bar plot displaying fraction of variants within rare, low-frequency and common allele frequency bins (as described for panel a) ordered by increasing proportion of common alleles (i.e. derived AF>1%) and excluding outliers (SJINF018, SJINF021, SJINF010 and SJINF008). Proportion of RNA-seq germline and somatic variants identified by overlap with remission/diagnostic WES data are displayed for comparison. (D) Crystal structure of the MutS lesion recognition complex (PDB CODE 2O8D^56^) MSH2:MSH6 mismatch repair proteins (MSH2: green, MSH6: blue) bound to ADP and G:T mismatch DNA (orange) showing the location of the amino acid change (NM_000251:exon9:c.G1465A:p.E489K). (E) Crystal structure of Pol k (green and blue representing different molecules) in complex with a bulky DNA adduct (orange) (PDB CODE 4U6P^28^) showing locations of the amino acid change (NM_016218:exon7:c.G893A:p. R298H).

We conclude that despite carrying thousands of somatic mutations, the coding-genome of ultra-mutated patient P337 shows similarities to canonical *KMT2A-R* iALL cases, vis-à-vis rare allele prevalence and representation of common human genetic variation. The LOH and somatic mutations are highly targeted to sites of known human variation and appear to effectively induce a “shuffling” of the germline allele repertoire.

### RAS-pathway and DNA-replication and repair gene mutations

Gene ontology analysis of the expressed non-synonymous somatic mutations (n=535) revealed over-representation of RAS-pathway genes (7/23 genes; 7-fold enriched; P=0.049; *RALGDS, RAC1, CHUK, PIK3R1, NFKB1, PLD1* and *PIK3CA)* and *“MLL* signature 1 genes” (33/369 genes; 2-fold enriched; P=0.046). Of the 535 non-synonymous expressed somatic mutations, 45 were predicted deleterious/damaging, including in the MMR gene *MSH2* (p.E489K). The *MSH2* somatic mutation localizes to a hot-spot, within the MMR ATPase domain in a region at the periphery of the DNA binding site, approximately 20 Å from the DNA molecule in the complex structure of full-length MSH2, a fragment of MSH6, ADP and DNA containing a G:T mispair ^56^. The mutation encodes a substitution of a negatively charged glutamate with a positively charged lysine (Figure 4D). There was a non-synonymous somatic mutation in another MMR gene, *MUTYH;* however, this site is a common human allele with a derived AF of 0.297 and was not called as damaged/deleterious by our prediction pipeline. There were no somatic mutations, LOH-sites nor rare/deleterious coding germline alleles detected in *POLE* or *POLD1* which are frequently co-mutated with *MSH2* in ultra-mutated tumours ^9^, ^50^ and in sporadic early-onset colorectal cancer ^42^. Among the 45 expressed and predicted damaging somatic mutations, there were two rare substitutions in DNA polymerase encoding genes: *POLG2* (p.D308V; ExAC AF 0.0000165), which encodes a mitochondrial DNA polymerase and *POLK* (p.R298H; ExAC AF 0.0003), which encodes an error prone replicative polymerase that catalyzes translesion DNA synthesis. The *POLK* mutation maps to a conserved impB/mucB/samB family domain (Pfam: PF00817) involved in UV-protection. The encoded substitution of arginine to histidine is located approximately 20 Å from the DNA molecule in the structure of a 526 amino acid N-terminal Pol K fragment complexed with DNA containing a bulky deoxyguanosine adduct induced by the carcinogen benzo[a]pyrene (Figure 4E) ^28^. Both wild-type and mutant residues are positively charged, however arginine is strongly basic (pI 11) while histidine is weakly basic (pI 7) and titratable at physiological pH ^52^. We additionally observed an expressed and predicted functional germline rare allele in *ATR*, a master regulator of the DNA damage response (p.L2076V; ExAC AF 0.0002).

### Global transcriptional similarities of the ultra-mutated and canonical infant leukemias

We next examined the transcriptional profile of the highly mutated *KMT2A-R* iALL specimen in comparison to a collection of patient leukemia samples and purified blood progenitor populations (Figure 5). As expected cell type was the major source of variation distinguishing normal hematopoietic progenitor, lymphoblastic leukemias and myeloid leukemias. We also observe clustering according to *KMT2A* translocation status and partner genes as reported previously ^51^. Replicate samples of the highly mutated *KMT2A-R* iALL specimen cluster with canonical counterparts demonstrating global transcriptional similarities (Figure 5).

**Figure 5:** Global transcriptional concordance of ultra-mutated infant leukemia. Multi-dimensional scaling of RNA-seq count data from acute leukemias and normal bone marrow progenitor populations displaying translocation partner gene by color and blood/leukemia type by symbol shape according to the legend. Labels show replicate measurements from the ultra-mutated diagnostic specimen, P337.

## Discussion

We report sequence analysis of an ultra-mutated acute lymphoblastic leukemia diagnosed in an infant at 82 days of age which, to our knowledge, represents the earliest onset of an ultra-mutant tumor recorded. The mutation rate observed was in range of cases reported previously in childhood cancers ^9^, ^50^. By integrating analysis of somatic mutations and germline alleles with a larger cohort of *KMT2A-R* iALL patients we found that the high somatic mutation rate does not generate large numbers of rare alleles compared to typical cases. Rather, we observe that the ultra-mutated *KMT2A-R* leukemic coding genome closely resembles the rare allele landscape of canonical cases. This apparent paradox is explained by the net effect of somatic gains and losses, both of which were targeted to rare and common alleles in proportions that off-set the depleted pool of germline alleles observed in the leukemic coding sequences. Our data indicate both the rare allele burden and transcriptional profile of the ultra-mutated leukemia were comparable to its canonical counterparts. Therefore, despite its radically different genetic route to leukemogenesis, the molecular profile of the ultra-mutated specimen seems to have converged towards that of canonical cases.

Most previously reported ultra-mutated childhood tumours have a combined MMR and proof reading polymerase deficiency ^9, 50^ - we detect evidence for the former *(MSH2* point mutation), but not the latter. However, we identify somatic mutations in two genes encoding DNA polymerases *(POLK* and *POLG2*) and an additional MMR gene (*MUTYH*). Notably, the *POLK* mutation is a rare allele, predicted functional and encodes Pol κ, a replicative error-prone translesion DNA polymerase that plays a role in bypassing unrepaired bases that otherwise block DNA replication ^29^, ^37^. For example, Pol κ preferentially incorporates dAMP at positions complementary to 7,8-dihydro-8-oxo-2’-deoxyguanosine adducts ^27^. As a corollary to these observations, *POLK* mutations have been found in prostate cancer with elevated G→A transitions ^57^. The largest fraction of ultra-mutations was attributed to Signature 1, associated with 5-methylcytosine deamination inducing C→T transitions, which accumulates in normal somatic cells with age over the human lifespan ^3^. This signature is found ubiquitously across tumour sub-types ^2^, and prominently in hypermutated early onset AML patients with germline *MBD4* mutations ^39^. LOH-sites (reference^-altemative) were enriched for Signature 5 which is also correlated with human aging. We observe enrichment of Signature 20 in LOH-sites and somatic alterations, consistent with an association with a MMR-defect ^2^. This signature has also been reported as enriched in tumours with germline-MMR and secondary mutation in the *POLD1* proof-reading gene ^2^. We also find enrichment of Signature 12 which arises from a poorly characterized underlying molecular process; however, the dominant base transitions, A→G and T→C are associated with DNA damage (e.g. by adducts) in normal and cancer tissues ^13^, ^36^. These observations suggest common mutagenic and repair deficits driving human genomic variation were operative in the cell of origin during pre-natal evolution of the ultra-mutated infant leukemia genome.

Our data indicate that most ultra-mutations are likely passengers coinciding with common alleles, consistent with previous findings ^38^; however, as proposed by Valentine and colleagues ^53^, a minority of rare alleles may contribute to infant leukemogenesis. Strikingly, LOH and somatic mutations occurred proportionally at rare and common polymorphic sites, such that there were no major changes in rare allele burden. Thus, the mutational process appears to precisely shuffle the patients’ constellation of inherited polymorphisms, with variations found in the wider human population. Due to the clonal nature of these events they appear to have occurred in rapid succession and may have been subject to positive selection resulting in the fixation of passengers ^19^. An alternative model is also possible in which the MSH2-mutation impairs DNA double-strand-break repair ^8^, ^54^ promoting *KMT2A*-rearrangement, but with neutral selection of ultra-mutations and loss of the mutator-phenotype. Both scenarios are consistent with the data presented here. Further studies are required to determine if combinations of somatic mutations and/or germline alleles promote *KMT2A-R* infant leukemogenesis and/or hypermutation.

The peculiar nature of this observation limits the scope of generalizable conclusions that can be drawn. However, by deeply investigating this unusual case we rule out technical artifacts (e.g. due to sample preparation or variant calling methodology) thereby demonstrating an unprecedented preponderance of ultra-mutated sites to overlap germline alleles proportional to their population frequencies. While this is the first demonstration (to our knowledge) of a hypermutated cancer with a somatic signature that precisely correlates with human germline polymorphic sites, similar signatures have been reported in both non-hypermutant^47^ and hypermutant^9^ sub-types. Since somatic mutation detection pipelines often remove sites of common human variation, concomitant germline and somatic hot-spots may currently be underestimated. The characterisation of additional tumours with a similarly skewed somatic signature is necessary to explore their clinical relevance and potential utility as biomarkers for personalised therapies.

## Materials and Methods

### Patient specimens, external sequencing data and controls

The study was approved by the Human Research Ethics Committee of the Princess Margaret Hospital for Children. The study cohort consisted of ten infants diagnosed at Princess Margaret Hospital for Children (Perth), of which we have previously reported RNA-seq data for five patients ^18^. Additional RNA-seq data was downloaded for 39 *KMT2A-R* iALL cases at diagnosis, six cases at relapse, six cases lacking the *KMT2A-R* (KMT2A-germline) and for pediatric *KMT2A-R* patients at diagnosis, comprising 10 AML cases, seven ALL cases and one undifferentiated case as reported previously by Andersson *et al*. ^5^ (accession EGAS00001000246). RNA-seq data comprising 28 *KMT2A-R* and 52 KMT2A-germline adult diagnosistic AML cases downloaded from the Short Read Archive (BioProject Umbrella Accession PRJNA278766^33^; Leucegene: AML sequencing). Normal human bone marrow transcriptomes were downloaded from the Short Read Archive (BioProject PRJNA284950) comprising 20 pooled sorted human hematopoietic progenitor populations ^10^. The full list of samples analysed in this study including those downloaded from external sources is summarised in Supplementary Table S1.

### Sample preparation and next-generation sequencing

RNA was isolated with TRIzol™ Reagent (Invitrogen Life Science Technologies, Carlsbad, CA, USA) and purified with RNeasy Purification Kits (Qiagen, Hilden, Germany). Genomic DNA was isolated with QIAamp DNA Mini Kit (Qiagen). Nucleic acid integrity and purity was assessed using the Agilent 2100 Bioanalyzer (Agilent Technologies, Santa Clara, CA). RNA-seq was performed at the Australian Genome Research Facility, Melbourne with 1 μg RNA as previously reported ^18^. WES was performed by the Beijing Genome Institute using SureSelect XT exon enrichment (Agilent Technologies) with 200 ng of remission DNA (v4 low input kit) or 1 μg of diagnostic DNA (v5). Diagnostic RNA and remission genomic DNA sequencing was performed on a HiSeq 2000 (Illumina, Inc., San Diego, CA, USA) instrument with 100bp paired end reads; diagnostic genomic DNA sequencing was performed on a HiSeq 2500 with 150bp paired end reads. Sequencing libraries were constructed using Illumina TruSeq kits (Illumina, Inc.). The sequencing data has been deposited in the European Genome Phenome Archive (ega-box-909).

### Bioinformatics

For gene-expression analysis, RNA-seq reads were mapped to hg19 with HISAT and gene-counts generated with HTseq ^4^. Data was normalized with RUV-seq ^45^ implementing factor analysis of negative control genes identified by fitting a linear model with cell type (hematopoietic sub-set or leukemia sub-type) as a covariate. voom! ^34^ was used to adjust for library size and generate counts per million (cpm) values. KMT2A-fusion split-reads were identified using RNA-seq data with *FusionFinder* ^23^ using default settings.

Single nucleotide variants (SNVs) were identified from RNA-seq data using the intersection of two methodologies: (1) the “Genome Analysis Toolkit (GATK) Best Practices workflow for SNP and indel calling on RNAseq data” and (2) *SNPiR* ^44^. WES data was processed based on the GATK Best Practices workflow. Variants were characterized using custom scripts that extract the coverage of sequenced bases from VCF and bam files, determining the overlap among samples, and calculating variant allele fractions (VAFs). Scripts used to execute variant detection software and analysis are available at https://github.com/cruicks/ultra-mutated-case under an MIT license updated at Github (https://github.com/cruicks/vcfbamCompare).

High confidence somatic mutations were identified using *VarScan2* (v2.3.9) *^31^* setting cut-offs of P-value<0.0005; coverage > 50X; VAF > 0.1 or VAF > 0.3 to enrich for clonal mutations. Variants were annotated with ANNOVAR ^55^ and with metadata from additional sources including seven functional/conservation algorithms (PolyPhen2 ^1^, Sift ^32^, MutationTaster ^49^, likelihood ratio test ^15^, GERP ^16^, PhyloP ^17^, and CADD ^30^) and databases of human genetic variation (Exome Aggregation Consortium [ExAC] ^35^, 1000 genomes, HapMap, dbSNP, Catalog of Somatic Mutations in Cancer (COSMIC)^22^ and ClinVar). Predicted functional alleles were assigned for variants called as damaged/deleterious by at least 4/7 algorithms.

Somatic mutational spectra were investigated using the R package, *DeconstructSigs* (v1.8.0) ^46^ calculating the fraction of somatic substitutions within the possible 96-trinucleotide contexts relative to the exome background. Copy number analysis was performed on WES data using *VarScan2* to calculate log2 depth ratio and GC-content and *Sequenza* (v2.1.2) ^21^ to identify chromosomal segments with copy loss. Gene ontology enrichment was performed using GREAT (v3.0.0) ^40^ bioinformatics tools. Mutations were inspected by mapping the genomic position to crystal structures in the Protein DataBank^7^ and analyzed with Pymol (v1.8.6.2). Sample distances displayed as a dendrogram were based on variant calls and calculated with SNPRelate (v3.6) ^58^.

## Acknowledgments

This work was supported by the Children’s Leukaemia and Cancer Research Foundation (Inc.) (project grants to URK, MNC, RSK; Fellowship to MNC); the Ian Potter Foundation (AG); Healy Foundation (AG); Telethon Kids Institute Blue Sky Research Grant (MNC, URK, RSK, JW); Telethon Kids Institute Research Focus Area Precision Medicine Working Group Funding (JW, MNC); NHMRC Career Development Fellowship 1066869 (JW); and Raine Medical Research Foundation Clinician Research Fellowship 2016–2018 (RSK). We thank Daniel MacArthur and Fengmei Zhao for assisting with exome sequence analysis; and researchers at The Pediatric Cancer Genome Project at St. Jude Children’s Research Hospital for provision of data. We thank all the patients, families and hospital staff for providing samples.

## List of abbreviations

Histone-lysine N-methyltransferase 2A (KMT2A), infant acute lymphoblastic leukemia (iALL), acute myeloid leukemia (AML), Histone-lysine N-methyltransferase 2A gene rearrangements (KMT2A-R), single nucleotide variants (SNVs), whole exome sequencing (WES), The Pediatric Cancer Genome Project (PCGP), Catalog of Somatic Mutations in Cancer (COSMIC), variant allele fraction (VAF), loss-of-heterozygosity (LOH), single nucleotide polymorphisms (SNPs), mismatch repair (MMR).

## Authors’ contributions

AG wrote computer code, performed bioinformatics, conceived analysis strategy, interpreted and analyzed data and edited the manuscript. AA and BF performed experiments. KWC, PK and CB performed additional data analysis and interpretation. RSK provided patient samples and clinical data, assisted with data interpretation and edited the manuscript. CHC, URK and JW acquired funding support, assisted with data interpretation and edited the manuscript. MNC performed bioinformatics, formulated research, conceived analysis strategy, acquired funding support, interpreted data, designed experiments and drafted the manuscript.

---

**Table 1:** Characterisation of somatic lesions in *KMT2A-R* patient samples.

| Patient ID | *KMT2A*-fusion | Reciprocal *KMT2A*-fusion | Diagnostic leukemia DNA | Remission DNA | Point mutations 0.1 > VAF<sup>1</sup> < 0.3 | Point mutations VAF > 0.3 | Indels VAF > 0.3 | LOH<sup>2</sup>-sites | Point mutations or hotspots with RNA-SEQ reads |
| --- | --- | --- | --- | --- | --- | --- | --- | --- | --- |
| P337 | *KMT2A-MLLT1* | Not detected | Exome | Exome | 198 | 5,054 | 260 | 6,555 | > 1,000 expressed somatic mutations |
| P401 | *KMT2A-MLLT3* | Not detected | Exome | Exome | 78 | 3 | 9 | 45 | No non-silent somatic mutations |
| P399 | *KMT2A-AFF1* | *AFF1-KMT2A* | Exome | Exome | 95 | 3 | 7 | 67 | No non-silent somatic mutations |
| P438 | *KMT2A-MLLT3* | Not detected | Exome | Exome | 74 | 4 | 9 | 100 | NRAS.p.G12S |
| P810 | *KMT2A-EPS15* | *EPS15-KMT2A* | Exome | Exome | 95 | 4 | 5 | 46 | PER3.p.M1006R, PER3.p.K1007E |
| P809 | *KMT2A-EPS15* | *EPS15-KMT2A* | Exome | Exome | 69 | 3 | 3 | 28 | PER3 M1006R, PER3.p.K1007E |
| P848 | *KMT2A-AFF1* | *AFF1-KMT2A* | no | Exome | NA | NA | NA | NA | KRAS.p.A146T |
| P287 | *KMT2A-AFF1* | *AFF1-KMT2A* | no | no | NA | NA | NA | NA | No COSMIC<sup>3</sup> hotspots |
| P706 | *KMT2A-MLLT1* | Not detected | no | Exome | NA | NA | NA | NA | KRAS.p.A146T |
| P272 | *KMT2A-AFF1* | Not detected | no | no | NA | NA | NA | NA | No COSMIC<sup>3</sup> hotspots |

<sup>1</sup>VAF: Variant allele fraction; <sup>2</sup>LOH: Loss-of-heterozygosity; <sup>3</sup>COSMIC: Catalogue of Somatic Mutations in Cancer

**Table 2:** Mean and standard deviation of expressed rare (Not in ExAC or <0.5% allele frequency), low-frequency (0.5–1% allele frequency) and common (>1% allele frequency) alleles in 39 *KMT2A-R* patient samples compared to ultra-mutated (P337) sample and its constituent expressed germline alleles and somatic mutations.

|  | **Not in**<br>**ExAC** | **<0.5% AF<sup>1</sup>**<br>**in ExAC** | **0.5-1% AF**<br>**in ExAC** | **>1% AF in**<br>**ExAC** |
| --- | --- | --- | --- | --- |
| *KMT2A-R* iALL | 42.3±10.4 | 212.1±32.6 | 76.8±14.7 | 7722.3±266.0 |
| P337 expressed | 38 | 174 | 89 | 7590 |
| P337 Germline | 17 | 88 | 47 | 6235 |
| P337 Somatic | 21 | 86 | 42 | 1355 |

<sup>1</sup>AF: allele frequency; <sup>2</sup>ExAC: Exome aggregation consortium

## Supporting information captions

**Supplementary Fig. S1.** Copy-number alterations and loss-of-heterozygosity overlap in typical and ultra-mutated *KMT2A-R* iALL. (a) Genomic copy number ratios of the ultra-mutated sample comparing diagnostic and remission WES data. (b) Proportion of reads mapping to different copy-number bins (up to 4). (c) Box and whisker plots (median with upper and lower quartiles) displaying the nearest distance of a variant/mutation to a homozygous variant in the diagnostic WES data for canonical samples (left) compared to the ultra-mutated sample (right) for VarScan2 germline, somatic and LOH SNV calls.

**Supplementary Table S1:** Summary of samples

**Supplementary Table S2:** Read evidence of RNA-seq variants that were not detected in matched WES data.

**Supplementary Table S3:** Comparison of RNA-seq reads from diagnosis sample with WES reads.

**Supplementary Table S4:** Comparison of RNA-seq reads from diagnosis sample with WES reads from diagnosis, remission and derived cell lines.

**Supplementary Table S5:** Annotation of somatic mutations and indels from ultra-mutated patient P337.

**Supplementary Table S6:** Summary of LOH-sites called in primary P337 and derived cell lines.

**Supplementary Table S7:** Summary of somatic SNVs called in primary P337 and derived cell lines.

**Supplementary Table S8:** Summary of quality filtered somatic SNVs and LOH-sites reads in primary P337 and derived cell lines.

